# Mechanisms of active regulation of biomolecular condensates

**DOI:** 10.1101/694406

**Authors:** Johannes Söding, David Zwicker, Salma Sohrabi-Jahromi, Marc Boehning, Jan Kirschbaum

## Abstract

Liquid-liquid phase separation is a key organizational principle in eukaryotic cells, on par with intracellular membranes. It allows cells to concentrate specific proteins into condensates, increasing reaction rates and achieving switch-like regulation. However, it is unclear how cells trigger condensate formation or dissolution and regulate their sizes. We predict from first principles two mechanisms of active regulation by post-translational modifications such as phosphorylation: In enrichment-inhibition, the regulating modifying enzyme enriches in condensates and the modifications of proteins inhibit their interactions. Stress granules, Cajal bodies, P granules, splicing speckles, and synapsin condensates obey this model. In localization-induction, condensates form around an immobilized modifying enzyme, whose modifications strengthen protein interactions. Spatially targeted condensates formed during transmembrane signaling, microtubule assembly, and actin polymerization conform to this model. The two models make testable predictions that can guide studies into the many emerging roles of biomolecular condensates.

---

Eukaryotic cells contain numerous types of membraneless organelles, which contain between a few and thousands of protein and RNA species that are highly enriched in comparison to the surrounding nucleoplasm or cytoplasm. These biomolecular condensates are held together by weak, multivalent and highly collaborative interactions, often between intrinsically disordered regions of their constituent proteins (Banani et al., 2017; Shin and Brangwynne, 2017).

In contrast to membrane-bound organelles, cells can regulate the formation and size of condensates by post-translational modifications of one or a few key proteins, most prominently by phosphorylation. The modifications usually lie within intrinsically disordered regions and modulate the strength of attractive interactions with other condensate components (Bah and Forman-Kay, 2016; Fung et al., 2018). Due to the highly cooperative nature of phase transitions, small changes in interaction strengths can result in the formation or dissolution of condensates, and this switch-like, dynamic nature makes them ideal for regulation.

For instance the nucleolus, Cajal bodies, splicing speckles, paraspeckles, and PML bodies in the nucleus and P-bodies in the cytoplasm have to be dissolved during mitosis and reformed afterwards to ensure a balanced distribution of their content to daughter cells (Rai et al., 2018; Dundr and Misteli, 2010). Stress granules form upon cellular stress and are dissolved when the stress ceases (Wippich et al., 2013).

Whereas these long-known, floating droplet or-ganelles are large enough to be visible using simpler light microscopic techniques, in the past years liquid-liquid phase separation has been implicated in multifarious processes in which – often sub-micrometer-sized – condensates are formed at particular sites in the cell: at sites of DNA repair foci (Altmeyer et al., 2015), Polycomb-mediated chromatin silencing (Tatavosian et al., 2019), transmembrane signalling (Banjade and Rosen, 2014; Case et al., 2019), microtubule formation (So et al., 2019; Huang et al., 2018; Hernández-Vega et al., 2017; Jiang et al., 2015), actin polymerization (Weirich et al., 2017), endocytosis (Bergeron-Sandoval et al., 2017; Miao et al., 2018), transcription (Cho et al., 2018; Sabari et al., 2018; Chong et al., 2018; Boehning et al., 2018), at presynaptic active zones (Wu et al., 2019; Zeng et al., 2018), and for RNP transport (Formicola et al., 2019; Alami et al., 2014). Such localized condensates form upon a local stimulus to recruit the required set of proteins and are dissolved once the job is done.

Cells do not only need to regulate the formation and dissolution of each type of condensate. They also need to tightly regulate their size and with it their numbers, to allow many condensates to form in different locations, for instance to activate genes at thousands of active promoters. Here, we propose two active mechanisms used by cells for these purposes.

## Phase separation and condensate size behaviour

To keep the model simple, we consider only one type of condensate protein. In the dilute regime below the saturation protein concentration *c*_out_, condensate droplets cannot form (Figure 1A). Above *c*_out_, in the phase separation regime, condensates can be stable.

**Figure 1:**
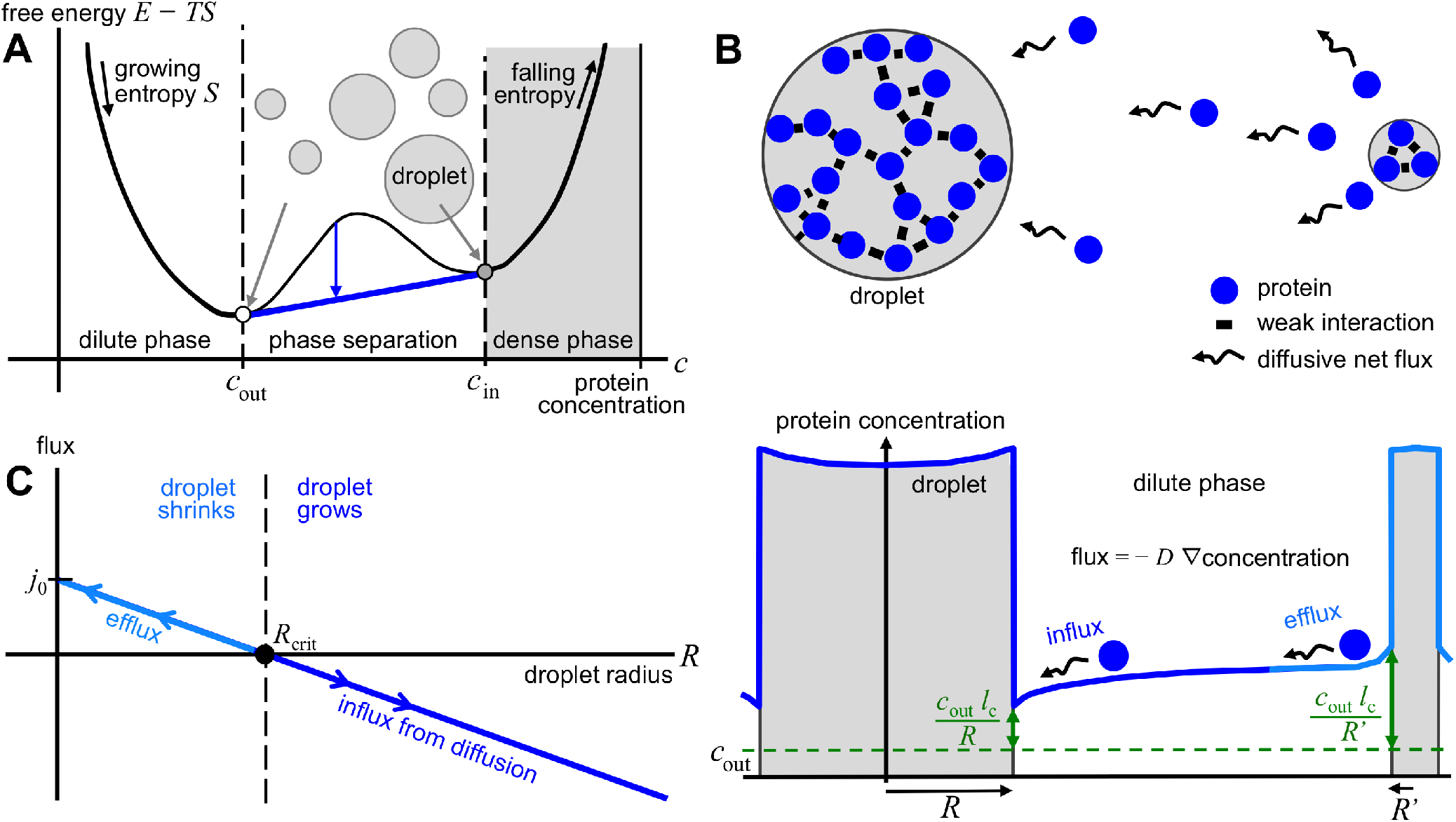
Phase separation and condensate droplet size behaviour. **A** When protein-protein and solvent-solvent interactions are more favorable than protein-solvent interactions, demixing into two phases can occur, a dilute phase with low protein concentration *c*_out_ and a dense phase with high concentration *c*_in_. This happens when the sum of free energies of the two phases is lower (tip of blue arrow) than the energy of the single phase (base of blue arrow). **B** *c*_out_ is the limiting concentration for infinite condensate droplet radius *R*. The concentration on the outside of a condensate of radius *R* is larger the smaller the condensate is (green double-headed arrows), as it cannot hold on to its proteins as well as large ones. This leads to a concentration gradient (∇concentration), which fuels a diffusive flux from small to large condensates (wiggly arrows). (*l*_c_ is a measure of interaction strength between proteins in comparison to the solvent.) **C** As a result, condensates below a radius *R*_crit_ will shrink and larger ones will grow.

However, in a passive system more than one condensate can not exist in equilibrium because larger condensates will grow at the expense of smaller ones (Figure 1B). To understand it, note that, in comparison to a large droplet, in a small droplet proteins on the surface have fewer favorable interactions with other droplet proteins due to its larger surface curvature. They are therefore more easily lost, resulting in a higher equilibrium concentration outside the droplet (For a quantitative derivation see Supplementary Material.) Due to this size dependence, the protein concentration decreases from small to large condensates, and the decrease generates a diffusive flux in the direction of steepest descent. Consequently, there exists a critical radius *R*_crit_ below which condensates will shrink while condensates above *R*_crit_ will grow (Figure 1C). The critical radius increases until a single large condensate survives, a phenomenon called coarsening (Hyman et al., 2014).

Here we show that, to actively regulate the formation and size of liquid droplet condensates, two generic mechanisms exist. An intracellular protein concentration maintained above saturation naturally leads to the enrichment-inhibition model, in which a droplet component-modifying enzyme such as a kinase inhibits favorable interactions and is enriched in condensates. A concentration maintained below saturation leads to the localization-induction model, in which the enzyme is localized or attached and induces favorable interactions. Although very simplified, these models probably capture the two essential mechanisms for regulating cellular condensates.

## The enrichment-inhibition model

Above the saturation concentration, a mechanism must exist that limits the size of larger condensates to allow for the coexistence of multiple condensates. This can be achieved if the loss of proteins from the condensate increases faster with condensate radius *R* than the gain by net diffusive influx. The influx is proportional to *R* − *R*_crit_ (Figure 1C, see Supplementary Material for a derivation). A loss that scales with condensate volume (4*π*/3)*R*^3^ would grow faster than *R* − *R*_crit_. Above a certain radius, the loss would surpass the influx, shrinking condensates that are too large and thereby resulting in a stable condensate size.

We propose the loss mechanism to be the modification of condensate proteins or RNA by an antagonistic regulating enzyme (orange) that is itself enriched in the condensate (Figure 2A). We use phosphorylation as an example, but the mechanisms work the same for other modifications (see below). The phosphorylation rate scales with the condensate volume if the concentration of unphosphorylated proteins (blue) is approximately constant in the condensate. In this model, unphosphorylated proteins as well as the kinases attract each other, while the phosphorylation weakens the interactions with other condensate proteins.

**Figure 2:**
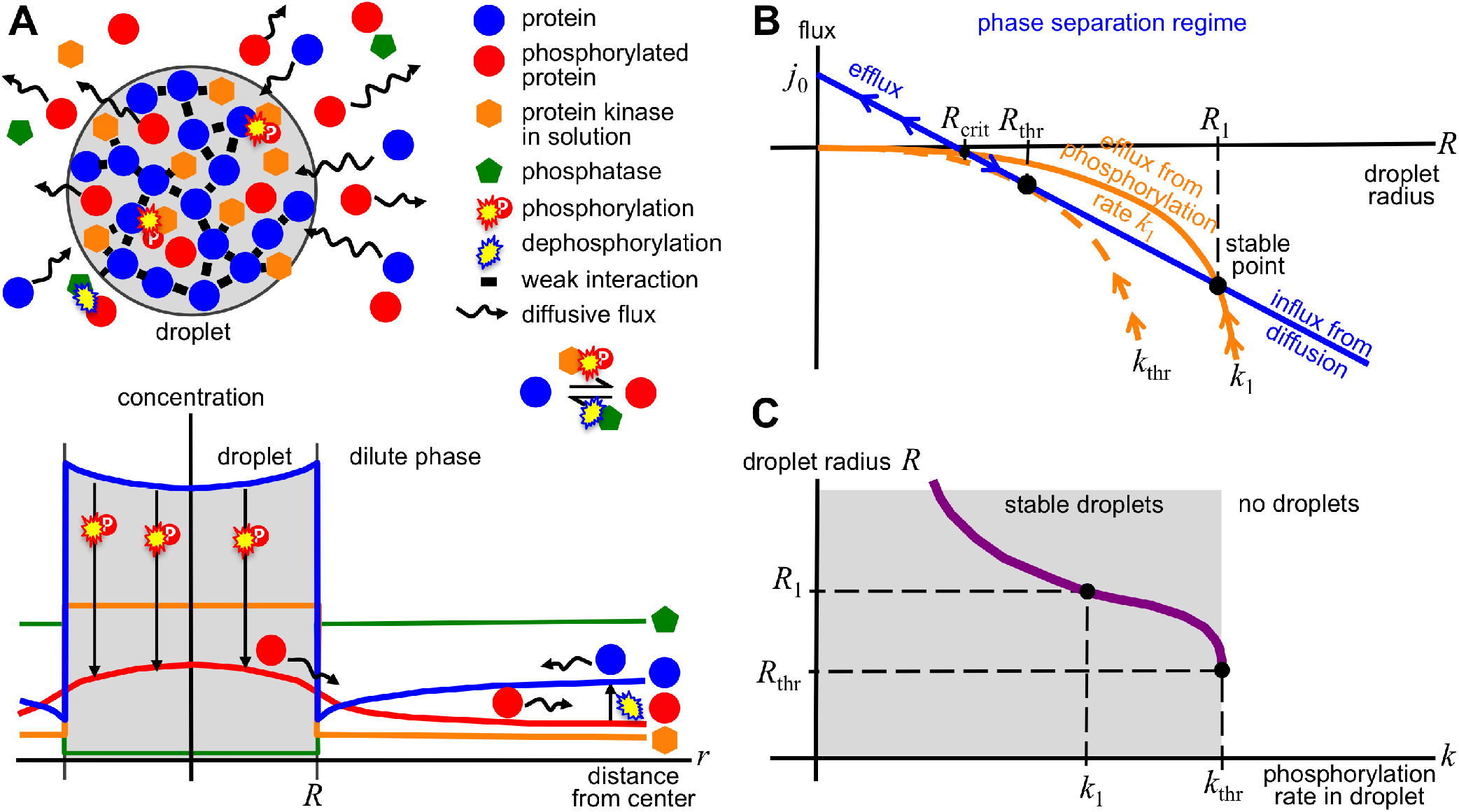
Enrichment-inhibition model. Condensate droplets can form when the concentration of unphosphorylated proteins is above the saturation concentration *c*_out_. Phosphorylation weakens protein interactions, so condensates can be dissolved by increasing kinase activity. **A** The unphosphorylated proteins (blue) and kinase (orange) get concentrated in the condensates via multivalent, attractive interactions. There, the kinase phosphorylates condensate proteins (red bursts). The non-interacting phosphorylated proteins (red) diffuse out of the condensate. Outside, they get dephosphorylated (blue burst). Unphosphorylated proteins diffuse back into the condensate along the gradient of concentration (blue), compensating the outward flux of phosphorylated proteins. **B** Without losses from phosphorylation, condensates in the phase separation regime grow by diffusive influx if their radius *R* is above a critical value *R*_crit_, whereas small condensates shrink (blue arrows). Because the influx grows linearly with condensate radius *R* whereas loss through phosphorylation grows with the condensate volume (4*π*/3)*R*^3^ (orange), a stable radius *R*_1_ results. **C** This radius depends on the phosphorylation rate *k* and shows a switch-like response.

Since the concentration of the unphosphorylated proteins is above saturation, the concentration decreases towards the condensate, leading to a net influx of unphosphorylated proteins (Figure 2A). This influx is compensated by the loss of proteins that get phosphorylated inside the condensate, which diffuse out along the negative concentration gradient. Outside, they are dephosphorylated by phosphatases (green), closing the circle of protein flux.

To avoid wasting energy by a short-circuited phosphorylation-dephosphorylation reaction, the phosphatase and kinase would best be concentrated in different phases. Therefore, we expect the phosphatase to be strongly depleted in the condensates.

Figure 2B shows that, for phosphorylation rates *k* below a certain threshold *k*_thr_, all condensates will grow or shrink to the same stable radius *R*, which is determined by *k*. The dependence of *R* on the phosphorylation rate *k* has a switch-like behaviour (Figure 2C): Above *k*_thr_, no condensates can exist.

It can seem counter-intuitive at first that the droplet-dissolving kinase enriches in the droplet. Yet, it is exactly this feature that allows the droplet growth to be self-limiting.

## The localization-induction model

Below the saturation concentration, no condensates can form. However, it is possible to *locally* push the concentration of proteins above saturation, in a small volume. This could be achieved if, in contrast to the previous model, the kinase acts agonistically and the phosphorylated proteins are attracted to each other through multivalent interactions, while the unphosphorylated proteins have little or no affinity for each other and other condensate proteins. With that assumption, the locally confined phosphorylation represents a source of condensate proteins that can compensate the diffusive loss that, below saturation, would otherwise cause the fast shrinkage and disappearance of any condensate.

Figure 3A illustrates the model. Phosphorylated proteins get generated at the site where the kinases are attached or bound, near the center of the condensate (Zwicker et al., 2018). From there, phosphorylated proteins diffuse out along the negative gradient of the concentration, which drops towards the outside. Outside of the condensate, the phosphorylated proteins get dephosphorylated by phosphatases. The dephosphorylated proteins diffuse in towards the center of the condensate to the point of lowest concentration, where the kinase activity depletes them, closing the circle of protein flux.

**Figure 3:**
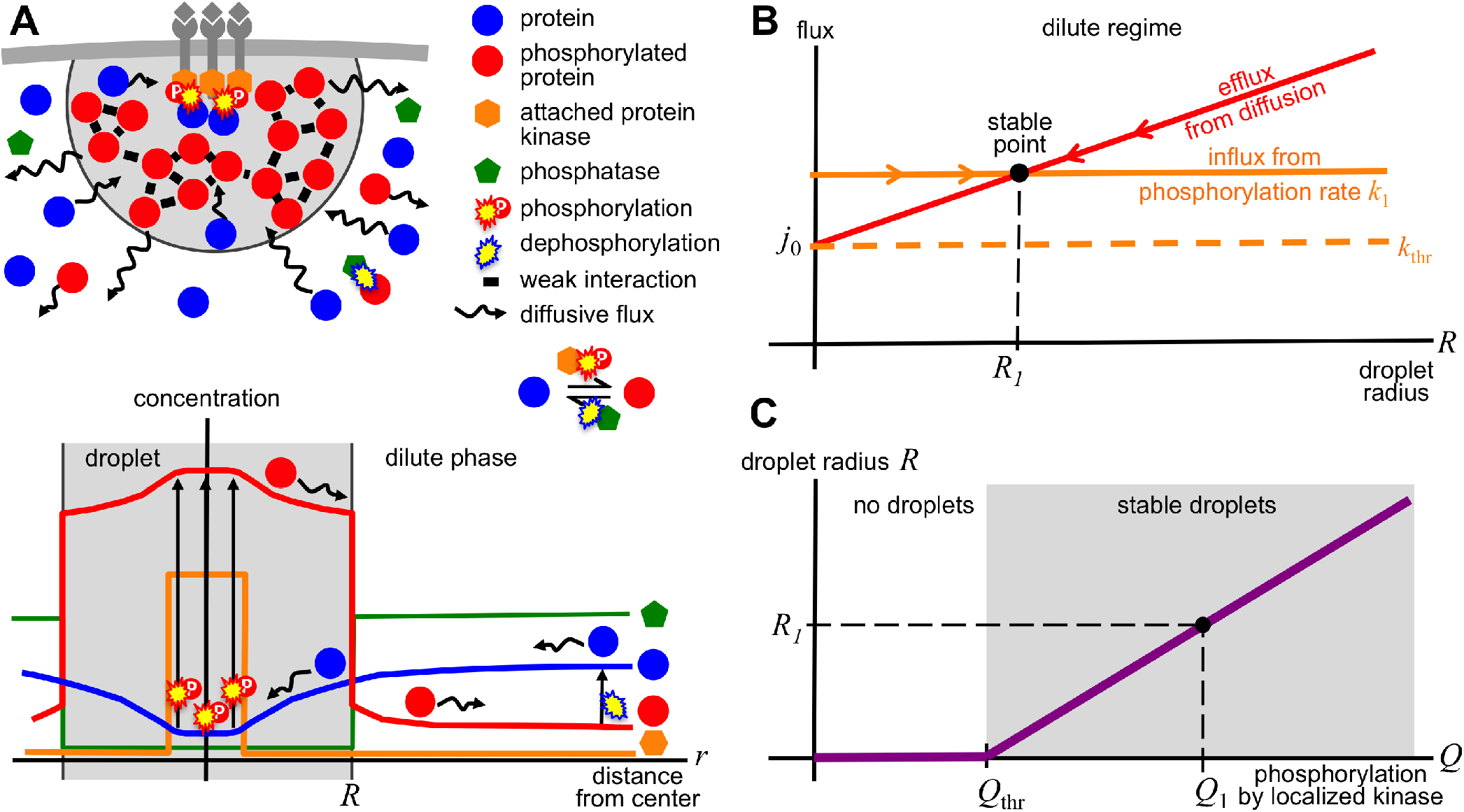
Localization-induction model. **A** Condensates can form when kinases (orange), bound or attached to a cellular structure such as a membrane, locally phosphorylate proteins (blue). This locally raises the concentration of the phosphorylated proteins (red), which can bind to each other through multivalent interactions, above the threshold for phase separation. Because the concentration of phosphorylated proteins outside the condensate is below saturation, the condensate loses phosphorylated proteins through diffusive flux (wiggly arrows). Outside, they get dephosphorylated (blue burst). Unphosphorylated proteins diffuse back into the condensate, compensating the outward flux of phosphorylated proteins. **B** Without kinase activity (orange), condensates in the dilute regime would shrink rapidly by diffusive efflux. Because the kinase activity supplies phosphorylated proteins at a constant rate *Q*_1_, a stable equilibrium is reached at radius *R*_1_. Below rate *Q*_thr_, no condensates can form. **C** Above *Q*_thr_, stable condensates form, whose radius depends linearly on *Q*.

As in the previous model, we expect the phosphatase to be strongly depleted in the condensates to minimize waste of ATP by premature dephosphorylation.

Figure 3B shows that, for phosphorylation rates *Q* below a certain threshold *Q*_thr_, no condensates can form because the efflux from a tiny drop is larger than the rate with which phosphorylated proteins are generated by the kinases. Above *Q*_thr_, the condensate radius *R* depends linearly on kinase activity (Figure 3C).

## Size regulation of transmembrane receptor clusters

Localization-induction, applied to phase transitions in two dimensions, can explain observations that at least some types of transmembrane receptors organize into dense clusters of fixed size upon binding to extracellular ligands (Case et al., 2019; Chamma et al., 2016; Thievessen et al., 2013; Rogacki et al., 2018).

Analogous to three dimensions, phase separation can occur in two dimensions above a certain saturation surface density, with the formation of clusters of high density surrounded by low-density regions (Banjade and Rosen, 2014; Case et al., 2019). However, analogous to the 3D case, large clusters would grow at the expense of small ones in an unregulated fashion.

In the Supplementary Material we describe a model that explains how cluster size might be regulated to a fixed size. Ligand-bound receptor kinases can cross-phosphorylate each other. Phoshorylation increases the affinity for binding each other. Stable cluster sizes are obtained from a balance of the diffusive influx, which in 2D depends only weakly on the cluster radius *R*, and a loss of receptors from the cluster through receptor dephosphorylation, which scales between *R* and *R*^2^.

## Regulation by other modifications

Regulation of condensate formation and size requires a signal input. This will mostly be the activity level of a kinase. However, other posttranslational modifications can take the role of phosphorylation. The posttranslational modification reaction, driven by ATP or some other high-energy compound, will supply the energy to maintain the net diffusive flux required for the size-stabilizing losses or gains.

Examples could be SUMOylation, which mediates multivalent interactions with proteins carrying SUMO interacting motifs, e.g. in PML bodies (Weidtkamp-Peters et al., 2008), poly-ADP-ribosylation, which induces condensate formation at DNA repair foci (Altmeyer et al., 2015), arginine dimethylation, which modulates condensate-forming propensities of proteins in RNA granules (Qamar et al., 2018; Ryan et al., 2018; Nott et al., 2015), lysine acetylation and methylation (Gibson et al., 2019), ubiquitination (Danieli and Martens, 2018), and RNA modifications (Drino and Schaefer, 2018). It is also likely that some condensates are regulated via the activity of localized or droplet-enriched phosphatases (Vallardi et al., 2019).

## Regulatory complexity

We envisage that the regulation of many biomolecular condensates will be more complex than the two simple models in two regards. First, condensates will usually be regulated by several kinases or posttranslational modifiers, whose effects on interactions within the condensate all combine. In Cajal bodies, for instance, two kinases regulate their formation but only one of them is localized to Cajal bodies (see below).

Second, some processes will be regulated in a multi-step fashion. For example in signalling cascades, a cascade of different types of condensate subdomains might form, each regulated by a different kinase and each depending on the previous one for the activation of its condensate-regulating kinase.

We surmise that, irrespective of the nature of regulatory complexity, the active regulation of condensates can be understood in terms of a combination of the two simple models.

## Related theoretical work

(Zwicker et al., 2015) and (Wurtz and Lee, 2018a) described how biochemically driven processes can be utilized for active size regulation of condensates. The enrichment-inhibition model is very similar to their models in that an energy-consuming reaction, e.g. phosphorylation, drives the transition between an interacting and a non-interacting protein species. In (Weber et al., 2019), the enrichment-inhibition model and the localization-induction model correspond to externally and internally maintained condensates, respectively. We clarified a crucial assumption, the differential enrichment of the kinase, phosphatase, or ATP (Patel et al., 2017) inside and outside of condensates for efficient size regulation. Without that assumption, the chemical potential of phosphorylated and unphosphorylated proteins would be almost constant in space, the net fluxes would be minimal and size regulation impossible (Supplemental Material).

## Evidence supporting enrichment-inhibition

We give five examples of biomolecular condensates that behave as expected from the enrichment-inhibition model: (1) Their key condensate protein(s) get phosphorylated by a kinase, (2) increased kinase activity dissolves the condensates, and (3) the kinase is enriched in the condensates. The model predicts the main phosphatase to be depleted in the condensates. This information appears to be mostly unavailable.

In the one-cell embryo of the worm C. elegans, RNAs and proteins form condensates called *P granules*. These localize to the posterior end of the cell and after cell division end up in the one cell that will give rise to the germ line. P granules are highly enriched for the intrinsically disordered MEG proteins. (1) They are phosphorylated by MBK-2 kinase and dephorphorylated by PPTR-1 phosphatase. (2) Phosphorylation of MEGs promotes granule disassembly and dephosphorylation promotes assembly. Furthermore, (3) MBK-2 localizes to P granules (Wang et al., 2014).

The vertebrate ortholog of MBK-2, DYRK3, plays as central role as dissolvase of several types of membraneless organelles during mitosis (Rai et al., 2018). This suggests that, as for P granules, DYRK3 is involved in the size control of many other types of condensates (Rai et al., 2018) by the enrichment-inhibition model.

*Stress granules (SGs)* are another example. (1) They are regulated by DYRK3 kinase (Wippich et al., 2013). However, since DYRK family kinases are constitutively active, it is unclear how the stress signal could be quickly relayed via DYRK3. (Wurtz and Lee, 2018b) proposed a plausible mechanism: Upon stress, ATP levels can fall by 50%, within the same time scale of SG formation (Hofmann et al., 2012). Also, ATP depletion alone is sufficient to induce SG formation. The reduction in DYRK3 activity (*k* in Figure 2C) by ATP depletion reduces the level of phosphorylation of its targets, (1) several of which are key SG proteins. (2) The concomitant increase in favorable interactions then triggers SG formation. Indeed, consistent with the enrichment-inhibition model, (3) DYRK3 localizes to SGs (Wippich et al., 2013).

*Splicing speckles* concentrate proteins involved in pre-mRNA splicing. These factors possess a terminal low-complexity RS region enriched for arginine and serine, which is required for the multivalent interactions within the speckles. (1) The CLK kinase phosphorylates the RS domains of splicing factors, and (2) phosphorylation by CLK promotes disassembly of splicing speckles (Kwon et al., 2014). Finally, (3) CLK possesses itself an RS domain that is required and sufficient for its enrichment within the speckles (Colwill et al., 1996).

*Cajal bodies* are nuclear condensates defined by the key protein coilin. (1 & 2) Hyper-phosphorylation of coilin by Cdk2/cyclin E or Vrk1 dissolves them. In contrast to Vrk1, (3) Cdk2/cyclin is strongly enriched in Cajal bodies (Liu et al., 2000; Cantarero et al., 2015).

Neurotransmitter-containing *synaptic vesicles (SV)* form dense clusters at synapses. (Milovanovic et al., 2018) discovered that synapsin, the major constituent of the matrix around SVs, formed condensates under physiological conditions. The condensates concentrated small lipid vesicles, explaining SV clustering at synapses. (1) CaMKII kinase phosphorylates synapsin and (2) its activity dissolved SV clusters in vivo in the presence of ATP. (3) Finally, CaMKII localized to the condensates as expected (Milovanovic et al., 2018).

## Evidence supporting localization-induction

The localization-induction model predicts three features: (1) The key condensate protein(s) get phosphorylated by a kinase, (2) whose phosphorylation activity promotes condensate formation, and (3) the kinase is targeted or attached to a specific cellular location. Here we discuss examples for cellular processes that appear to be actively regulated by localization-induction.

*T-cell signal transduction* exemplifies localization-induction in two dimensions. The T cell receptor kinase gets phosphorylated and can then recruit and bind the membrane-bound ZAP70 kinase, which thereby gets localized (3). The phosphorylation of T cell receptors also turns on the activity of its kinase domain, which phosphorylates and thereby activates the bound ZAP70 kinase. (1) Activated ZAP70 phosphorylates the key membrane-bound protein LAT. (2) Phosphorylation enables favorable interactions with several other proteins, with which LAT then forms a quasi two-dimensional condensate at the inner plasma membrane surface. The condensate excludes the LAT-dephosphorylating phosphatase CD45, but recruits the machinery for actin assembly, which can form condensates of its own to induce cytoskeletal changes (Su et al., 2016).

Many transmembrane signalling processes start by the formation of transmembrane receptor clusters. Case and colleagues argue that many other transmembrane signalling processes (using glycosylated receptors, immune receptors, cell adhesion receptors, Wnt receptors, and receptor tyrosine kinases) possess features that suggest they too involve liquid-liquid phase separation (Case et al., 2019).

Another example for localization-induction is the *assembly of microtubules*. Metaphase centrosomes consist of a core structure called the centriole pair, surrounded by a condensed phase, the pericentriolar matrix (Conduit et al., 2015). (1) This condensate forms by phase separation of Cnn in Drosophila (Raff, 2019) and enhances the nucleation of microtubules during mitosis. (2) Condensate formation depends on phosphorylation of Cnn by PLK-1 kinase (3) which is concentrated at the centrioles (Fu and Glover, 2012; Zwicker et al., 2014). In accord with the localization-induction model (Figure 3C), the phosphorylation rate determines centrosome size (Conduit et al., 2014).

Microtubule growth and kinetochore loading are also promoted by localized BuGZ condensates, which enhance tubulin concentrations (Jiang et al., 2015). They are also able to recruit Bub3, which facilitates chromosome alignment through binding to kinetochores (Jiang et al., 2014; Toledo et al., 2014). In neurons, (1) tau condensates could nucleate localized microtubule bundles and enhance their growth (Hernández-Vega et al., 2017). (2) As expected, tau phosphorylation enhances condensate formation (Ambadipudi et al., 2017).

Finally, we speculate that the localization-induction mechanism could contribute to *transcription regulation*. Condensates can form at transcription initiation sites, recruiting RNA polymerase II and resulting in transcription (Boehning et al., 2018; Cho et al., 2018; Sabari et al., 2018; Chong et al., 2018). The location and size of these condensates needs to be tightly controlled to selectively allow for transcription at thousands of genomic regions.

## Other mechanisms of size control

On longer timescales, condensate formation is certainly controlled by modulating concentrations of key droplet components using transcriptional or translational regulation. Protein aggregation appears to be regulated by an active de-aggregation mechanism that only kicks in at large aggregate sizes, much above the critical radius (Narayanan et al., 2019). The two simple models might also not help to understand the regulation of the highly complex geometries of passive and active phases of chromatin (Gibson et al., 2019; Hilbert et al., 2018) by a multitude of localized, sequence-specific binding events that modulate interaction strengths through modifications to histone tails.

In centrosomes of C. elegans, size is limited by the exhaustion of centrosome material (Decker et al., 2011; Zwicker et al., 2014). Similarly, exhaustion of material might explain the size control of condensates formed around cytosolic dsDNA to launch an anti-viral immune response (Du and Chen, 2018). Also the liquid phase that catches diffusing mRNPs whose encoded protein is destined for insertion into the membrane at the rough endoplasmatic reticulum might simply be size-controlled by exhaustion of material (Ma and Mayr, 2018). In other cases, sizes of condensates are also passively controlled (Feric and Brangwynne, 2013; Hyman et al., 2014).

## Missing kinase specificity

The enrichment-inhibition model offers an explanation for the relatively low kinase specificity that is frequently observed in in-vitro phosphorylation experiments and often deviates from the kinase specificity in-vivo (Miller and Turk, 2018). Kinases might attain most of their specificity not from their catalytic domain but from their enrichment in specific types of biocondensates. The latter is likely to be mostly determined by their disordered regions and peptide binding modules. This hypothesis is supported by the high fraction of disordered regions in cyclins of cyclin-dependend kinases (CDKs) and in non-CDK kinases.

## Conclusion

An important advantage of active regulation of biocondensates is the ability to switch processes or biochemical reactions on or off in response to a signal composed of only a few molecules, such as a DNA double strand break. The extremely cooperative behaviour of phase transitions (even for very small condensates) explains how such weak signals can get translated into the formation of a condensate, which then recruits all components necessary to react to the signal. To understand the magnitude by which condensates can accelerate reaction kinetics (Stroberg and Schnell, 2018), assume that *n* proteins are required to form an oligomeric complex. If each component is enriched 10-fold within the condensate, by mass action the oligomerization rate would be ~ 10^*n*^-fold increased.

Another advantage of active regulation is the thresholding behavior it produces (Figures 2C and 3C), as this can suppresses low-intensity noise and improve the robustness of cellular decisions.

We propose two generic mechanisms of active regulation, one applying to above-saturation concentrations of condensate components and the other to below-saturation concentrations. We discussed many examples illustrating the use of these mechanisms by cells in various pathways.

The models make quantitative, experimentally testable predictions about the dependence of formation and size of condensates on kinase activity and component concentrations (Supplementary Materials).

The two mechanisms supply unifying principles to the often seemingly incoherent behaviour of various types of biomolecular condensates. The enrichment-inhibition model predicts condensates floating in the cytoplasm or nucleoplasm, such as most membraneless organelles, to be size-regulated by an antagonistic kinase enriched in the condensates. The localization-inhibition model predicts spatially and temporally localized condensates such as those formed at sites of DNA repair foci to be regulated by an agonistic postranslationally modifying enzyme at their center.

Notably, it predicts these condensates to be stable even at very small radii (Figure 3C), at which the characteristic properties of phase-separation are hard to observe by microscopy (Alberti et al., 2019). It is attractive to speculate that many previously detected protein clusters or foci, such as the transcriptional minihubs and hubs observed in (Chong et al., 2018), might actually be tiny, bona-fide phase-separated liquid droplet condensates. The functional relevance of condensates versus non-phase-separated protein aggregations is the strong cooperativity in phase separatation. It might, for example, be the cause of the high cooperativity often observed in transcriptional regulation (Park et al., 2019; Hnisz et al., 2017).

It is becoming clear that liquid-liquid phase separation is a fundamental concept underlying most aspects of cellular regulation in eukaryotes. We hope the presented models will help to guide experiments elucidating the functions of biocondensates and to understand their manifold roles in human diseases (Wang and Zhang, 2019).

## Acknowledgements

We thank Matthew Grieshop for discussions and Klaus Förstemann (LMU), Michael Satteler, (TUM), Axel Imhof (LMU), Melina Schuh (MPI-BPC), and Ralf Krätzner (UMG) for feedback to the manuscript. JS and SSJ acknowledge support by grants SPP1935 and SPP2191 of the Deutsche Forschungsgemeinschaft.

## Author Contributions

JS and DZ initiated the study. JS designed and prepared main figures. JS wrote manuscript with input from all authors. SSJ drafted introduction and evidence supporting localization-induction. MB drafted evidence supporting enrichment-inhibition. DZ and JK contributed to theoretical modelling.

## Supplementary Material

### Condensate droplet growth and shrinkage in a passive, phase-separating system

#### Protein concentration *c*_*R*_(*R*) just outside a condensate of radius *R*

We derive here the formula for the protein concentration just outside a condensate of radius *R* (Figure 4A),

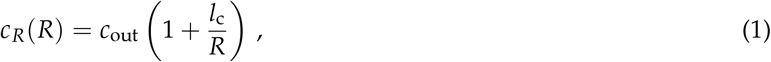

where *c*_out_ is the concentration outside a condensate droplet of infinite radius *R* ⟶ ∞ and *l*_c_ the capillary length, a measure of the strength of interaction between the proteins.

**Figure 4:**
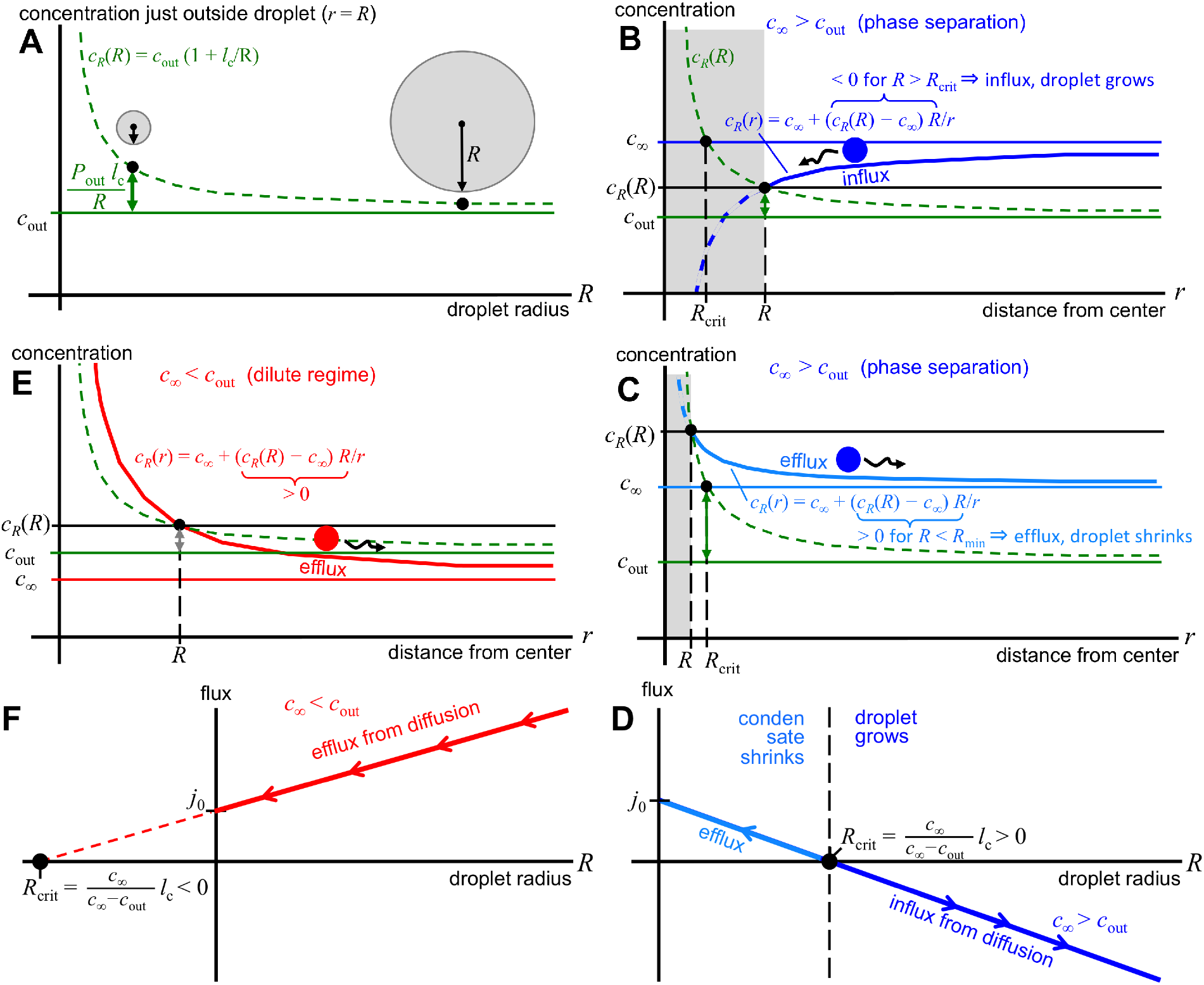
Concentration and flux around liquid droplet condensates in a passive s ystem. **A** The protein concentration just outside a condensate of radius *R* is higher by *c*_out_*l*_c_/*R* than the concentration *c*_out_ outside an infinitely extended phase (*R* ⟶ ∞). **B** Protein concentration around a condensate of radius *R* if the concentration *c*_∞_ at distance *r* ⟶ ∞ is larger than *c*_out_ and condensates above the critical radius, *R* > *R*_crit_. **C** Protein concentration for the case *c*_∞_ > *c*_out_ and condensates with *R* < *R*_crit_. **D** Net diffusive flux out of condensates of size *R* for *c*_∞_ > *c*_out_. **E** Protein concentration if *c*_∞_ < *c*_out_. **F** Net diffusive flux out of condensates of size *R* for *c*_∞_ < *c*_out_.

For a condensate of radius *R* we call the protein concentration just on its outside *c*_*R*_(*R*) and the concentration inside *c*_in_. We imagine to transfer from just the outside to the inside of the condensate a single protein together with just the right amount of solvent that this tiny volume *dV* has the same concentration as the inside of the droplet. That means 1 = *dV c*_in_, or 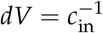. The radius of the condensate will increase from *R* to *R* + *dR*. *dR* can be obtained by solving the equation for the conservation of volume (4*π*/3)(*R* + *dR*)^3^ − (4*π*/3)*R*^3^ = *dV*, yielding

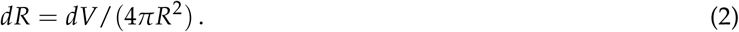

The change in free energy for the transfer must be zero at equilibrium between the two phases. The change of free energy is the change in interaction energy Δ*E* plus the change in chemical potential Δ*μ*, which accounts for the entropic effects:

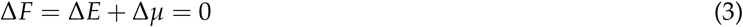

We define −*ε* as the change in interaction energy of the protein when transferred from the outside of an infinitely extended phase (*R* ⟶ ∞) to the inside. The change is slightly lower for a finite-sized condensate or radius *R*, due to surface tension:

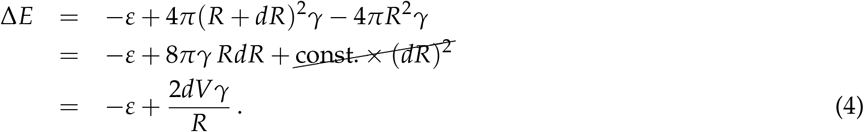

where the term 2*dVγ*/*R* = 2*γ*/(*c*_in_ *R*) describes the work done against the surface tension *γ* of the condensate. The change in chemical potential is

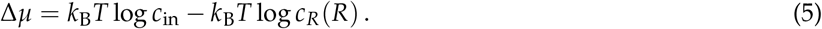

Putting everything together, we obtain

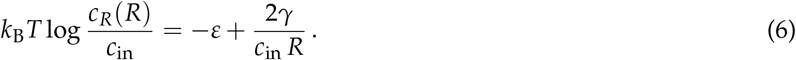

Defining the capillary length,

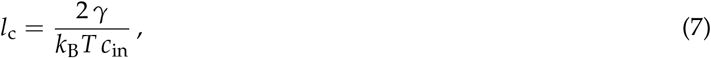

we get

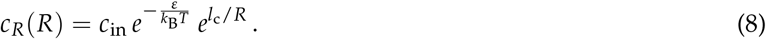

By setting *R* to ∞, we obtain 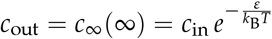 and therefore

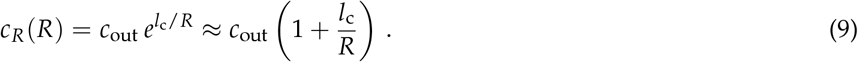

The approximation is accurate for *l*_c_/*R* ≪ 1. The finite-*R* correction term *c*_out_*l*_c_/*R* is called Laplace pressure in physics.

#### Concentration *c*_*R*_(*r*) around a condensate of radius *R* and net flux out of condensate

We imagine that only a single condensate of radius *R* floats in the dilute phase of infinite extension, and the protein concentration at infinite distance *r* from the condensate center is *c*_∞_. For symmetry reasons, the protein concentration will be the same everywhere on a spherical shell of radius *r* around the condensate, so we can write it as a *c_R_*(*r*).

From statistical physics we know that a concentration gradient ∇*p* causes a net diffusive flux density *j* in the opposite direction of the gradient, measured in particles per second per area through which the particles diffuse: **j**= − *D*∇*p*. The proportionality constant is the diffusion coefficient *D*. Applied to our system, the flux density through a spherical shell of radius *r* around the condensate is

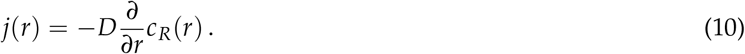

The total flux integrated over the entire shell surface 4*πr*^2^ is therefore 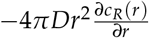. At equilibrium concentrations, the flux through each sphere of radius *r* > *R* must be constant, since no proteins can be created or lost between shells:

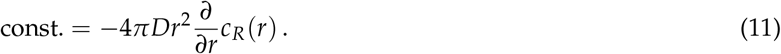

(The assumption of constant flux holds only approximately, for low rates of (de-)phosphorylation, in the active systems discussed in Figs 2 and 3.) This differential equation is solved by *c*_*R*_(*r*) = *α* + *β*/*r*. To obtain *α* and *β*, we use the boundary conditions at *r* = *R* and *r* = ∞,

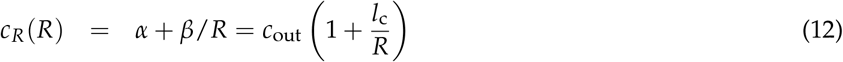

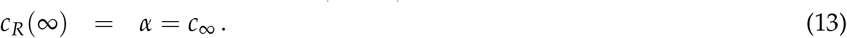

From this it follows that *β* = (*c*_*R*_(*R*) − *p*(∞))*R* and

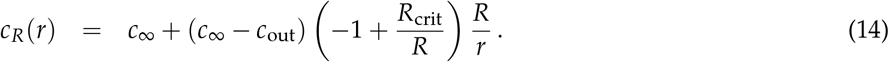

with

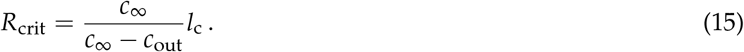

The total flux of proteins leaving the condensate is

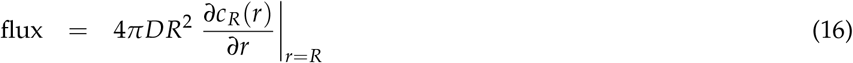

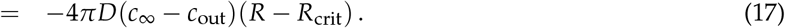

This result is plotted in Figure 4D for the case *c*_∞_ > *c*_out_. In that case, for condensates above the critical radius, *R* > *R*_crit_, the concentration decreases towards the condensate and the net flux flows towards the condensate (Figure 4B), making the condensate grow. For condensates below the critical radius, *R* < *R*_crit_, the concentration increases towards the condensate and the net flux flows away from the condensate (Figure 4C), making the condensate shrink. For the case *c*_∞_ > *c*_out_ (Figures 4E,F), the concentration increases towards the condensates for any radius, and the condensate flux will be directed outwards, shrinking the condensate. The critical radius for this case is negative and the flux depends on the radius as shown in Figure 4F.

#### Threshold radius *R*_thr_ and condensate radius *R*(*k*) for the enrichment-inhibition model

For the enrichment-inhibition model, we can determine the threshold radius *R*_thr_ of by solving *R* under the constraints that

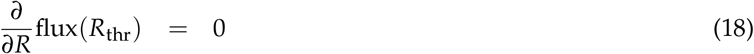

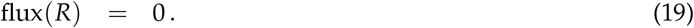

We insert the total flux

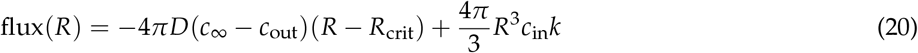

and obtain

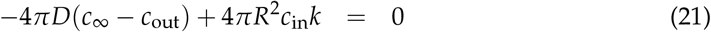

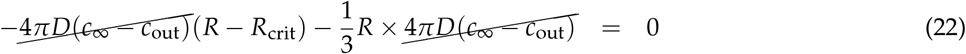

and hence

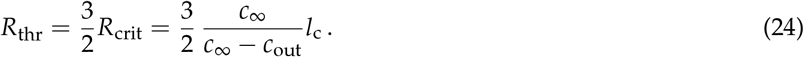

To find the condensate radius *R*(*k*) for a given phosphorylation rate per volume, *k*, with

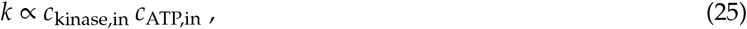

one can simply solve the cubic equation flux(*R*) = 0, which has an analytical solution (Cardano formula).

#### Condensate radius *R*(*Q*) for the localization-induction model

The radius *R*(*Q*) follows from the zero total flux condition

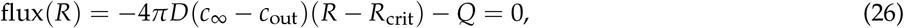

where *Q* is the total amount of phosphorylated proteins per time supplied to the condensate by the localized kinase. This yields, for *c*_∞_ < *c*_out_

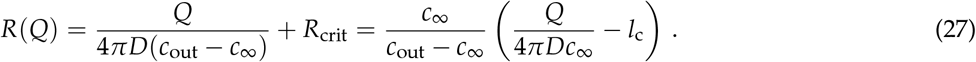

### Size regulation of transmembrane receptor clusters

Many transmembrane receptors organize into dense, supramolecular clusters of a certain preferred size upon binding to extracellular ligands (Case et al., 2019; Chamma et al., 2016; Thievessen et al., 2013; Rogacki et al., 2018). A fixed cluster size can be advantageous: Clusters need to be large enough for the kinase activity of the entire cluster to surpass the threshold rate *k*_thr_ for forming a localized liquid droplet at the membrane (Figure 3C). On the other hand, the uncontrolled size of receptor clusters would lead to an uncontrolled size of the localized droplets inside. Also, outsized clusters would deplete receptors and suppress signal transduction elsewhere.

Analogous to three dimensions, phase separation can occur also in two dimensions above a certain saturation surface density (Banjade and Rosen, 2014; Case et al., 2019), with the formation of clusters of high density surrounded by low-density regions. If ligand-bound receptors interacted while unbound did not, or more weakly, increasing the ligand concentration and thus the fraction of ligand-bound receptors would increase the surface density of ligand-bound, interacting receptors. When that density is pushed above the 2D saturation threshold, clusters would form. The mechanism would nicely explains the sharp onset of cluster formation upon a small change of ligand density. However, analogous to the 3D case, coarsening would lead to the growth of large clusters and shrinkage of small ones, not to clusters of fixed sizes.

Biophysical models can in principle explain the formation of clusters of fixed size by an interplay of short-range attraction between receptors and long-range repulsion (Case et al., 2019; Stradner et al., 2004) (just like the attractive, short-range strong nuclear force and the repulsive, long-range electrostatic force between protons in atomic nuclei explains the “valley of stability”). However, this model requires weak electrostatic screening (Case et al., 2019; Stradner et al., 2004), more precisely, the Debye length for the exponential decay of electric fields around charges must be at least on the order of the cluster size, whereas in fact in cells the Debye length is probably below a nanometer (Spitzer and Poolman, 2005).

The following two-dimensional version of the localization-induction model might explain how the cluster sizes can be regulated. Transmembrane receptors can exist in three states, unbound, bound, and phosphorylated. Ligand-bound receptors can cross-phosphorylate other receptors, unbound ones cannot. Active receptors get inactivated by dephosphorylation by a phosphatase with a certain rate. A stable cluster size should be achievable under the assumption that phosphorylated receptors bind to each other with higher affinity than unphosphorylated ones. Importantly, this assumption, which requires phosphorylation as an active, driven process, makes the ligand off-rate independent of whether the receptor is part of a cluster or not.

When a receptor gets dephosphorylated, the receptor can be lost from the cluster. If the mean net distance that unphosphorylated receptors travels by diffusion until it gets rephosphorylated is large in comparison to the cluster size, most receptor dephosphorylations will result in the loss or the receptor from the cluster, yielding a loss rate proportional to the number of receptors in the cluster times the phosphorylation rate, hence proportional to its radius squared, *R*^2^. If the mean net distance travelled is much smaller than the droplet radius *R*, only a fraction of receptors dephosphorylated near to the edge of the cluster will get lost from the cluster. In this case, the loss rate is proportional the circumference of the cluster, 2*πR*.

The loss is compensated by the influx of inactive receptors, which will be phosphorylated through contact with ligand-bound receptors in the cluster. The influx can be shown to roughly be proportional to (log *d*/*R*)^−1^, where *d* = (*πρ*)^−1/2^ is the root mean squared distance between clusters and *ρ* is the number of clusters of size *R* per cell surface area. The balance between the influx dropping with increasing *R* and the loss growing with *R* or *R*^2^ can then yield in a stable cluster size *R*.

### Enrichment of kinase, phosphatase or ATP is necessary for size stabilization

In simple systems without chemical reactions, biochemical condensates are regions enriched in droplet material that are in equilibrium with the surrounding solvent such that the chemical potentials are constant in space (*μ_i_* = const.); see Fig. 5A. Consequently, no spatial fluxes (which would be driven by gradients in chemical potential) exist. Because of surface tension, large droplets are energetically favorable, so that the equilibrium state is a single droplet whose size is determined by the total amount of phase separating material in the system.

**Figure 5:**
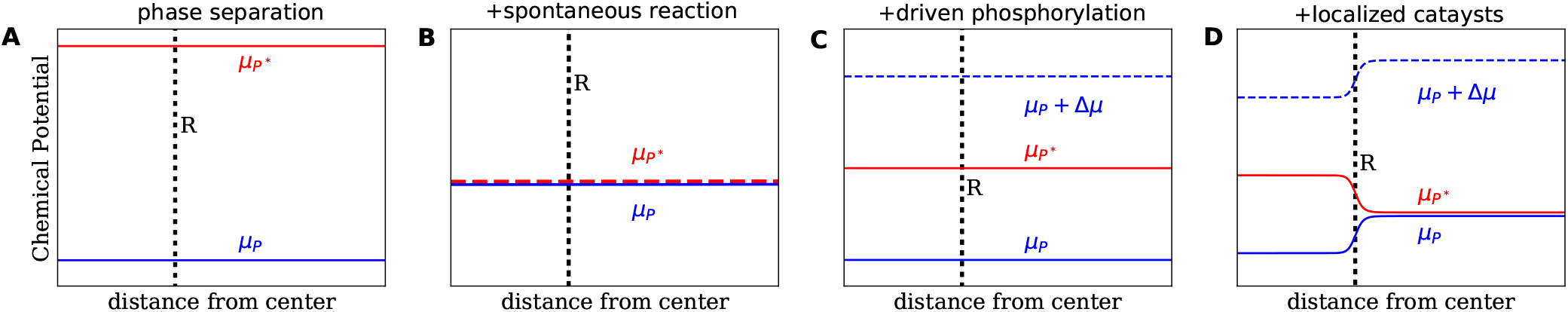
Schematic of the chemical potential of the unphosphorylated protein *P* (blue lines) and the phosphorylated protein *P\** (red lines) for different reaction schemes. **A** In the case of passive phase separation, the chemical potentials equilibrate independently. Consequently, gradients in chemical potential are absent and diffusive fluxes do not exist. **B** Adding spontaneous (de-)phosphorylation (*P* ⇌ *P\**) implies the chemical equilibrium *μ*_*P*_ = *μ*_*P*\*_. **C** Additionally driving the phosphorylation by ATP (*P* + ATP *P*\* + ADP) keeps the system away from equilibrium if ATP levels are maintained. However, if the reactions proceed equally in both phases phosphorylation and dephosphorylation can be balanced locally. Consequently, spatial fluxes do not exist although the energy Δ*μ* = *μ*_ATP_ − *μ*_ADP_ supplied by ATP is consumed. **D** Biasing the reactions, e.g. by localizing kinases and phosphatases in different phases, induces spatial fluxes and allows for size control.

If the droplet material can spontaneously transition between two states (e.g., between a phosphorylated state *P*\* and a dephosphorylated state *P*) the associated chemical potentials will equilibrate at every point in space (*μ*_*P*\*_ = *μ*_*P*_). In this case the reaction will change the relative amount of *P* and *P\** compared to the passive case, but after this chemical equilibration, the system is passive and no fluxes exist; see Fig. 5B.

Chemical fluxes can be sustained by an external energy input, e.g. by using ATP to phosphorylate *P* continuously. Since the metabolism of the cell maintains the chemical potentials of ATP, ADP, and phosphate constant, the conversion *P* ⇌ *P*\* with and without ATP cannot equilibrate at the same time. However, if these reactions proceed at the same rate everywhere, the phosphorylation driven by ATP will eventually be balanced by the dephosphorylation not involving ATP; see Fig. 5C. Consequently, although ATP is used continuously, there are still no spatial fluxes and the condensates behave qualitatively similar to the passive ones.

Kinases and phosphatases act as catalysts for the (de-)phosphorylation reaction, changing the reaction rates but not the equilibrium states. However, if the catalysts co-localize with the proteins, the reaction is biased towards different states inside and outside the condensate. This local bias leads to a chemical potential difference between the condensate and the surrounding, which drives spatial fluxes and can lead to a suppression of coarsening; see Fig. 5D. A strong segregation of kinases and phosphatases in different phases leads to strong spatial fluxes and thereby to a high conversion efficiency between the two species.

